# Reward and punishment enhance motor adaptation in stroke

**DOI:** 10.1101/106377

**Authors:** Graziella Quattrocchi, Richard Greenwood, John C Rothwell, Joseph M Galea, Sven Bestmann

## Abstract

The effects of motor learning, such as motor adaptation, in stroke rehabilitation are often transient, thus mandating approaches that enhance the amount of learning and retention. Previously, we showed in young individuals that reward-and punishment-feedback have dissociable effects on motor adaptation, with punishment improving adaptation and reward enhancing retention. If these findings were able to generalise to stroke patients, they would provide a way to optimize motor learning in these patients. Therefore, we tested this in 45 chronic stroke patients allocated in three groups. Patients performed reaching movements with their paretic arm with a robotic manipulandum. After training (day 1), day 2 involved adapting to a novel force-field. During this adaptation phase, patients received performance-based feedback according to the group they were allocated: reward, punishment or no feedback (neutral). On day 3, patients readapted to the force-field but all groups now received neutral feedback. All patients adapted, with reward and punishment groups displaying greater adaptation and readaptation than the neutral group, irrespective of demographic, cognitive or functional differences. Remarkably, the reward and punishment groups adapted to similar degree as healthy controls. Finally, the reward group showed greater retention. This study provides, for the first time, evidence that reward and punishment can enhance motor adaptation in stroke patients. Further research on reinforcement-based motor learning regimes is warranted to translate these promising results into clinical practice and improve motor rehabilitation outcomes in stroke patients.

## INTRODUCTION

Upper limb (UL) paresis is a common post-stroke outcome. Although rehabilitation can lead to improvements, the benefits are often inconsistent[1]. Principles of motor learning may offer ways to increase the efficacy of rehabilitation. This is underpinned by two assumptions: these principles apply to motor recovery, and patients retain the ability to learn[2]. However, only a few studies have investigated the effect of stroke on motor learning, with mixed outcomes[3–8], and interventions based on motor learning principles are often no more effective than conventional rehabilitation[9].

Motor adaptation is a specific form of learning that refers to gradual error reduction in response to a novel perturbation. Stroke patients retain the ability to adapt, even if at a slower rate than healthy individuals[4,6,7]. In particular, error-enhancing force-fields, i.e. magnifying movement error, appear more beneficial than error-reducing ones, as they lead to after-effects which compensate for the original error[4,10]. We focus on motor adaptation as a model process to investigate learning in a standardized way within a single session.

Reward and punishment-based feedback are candidate mechanisms to optimize motor adaptation[11–13]. In young healthy participants, punishment was associated with faster adaptation, and reward with greater retention[13]. These results point to dissociable effects of reward and punishment in motor adaptation. If these findings generalised to patients, they would provide a principled way for enhancing motor adaptation and retention in stroke survivors. This would be in line with previous research showing the benefits of rewards during ankle boot training in stroke survivors [14]. Using a force-field adaptation reaching task, we tested this in 45 chronic stroke patients. We show that reward-and punishment-based feedback enhance stroke patients’ motor adaptation, and that reward increases the retention of the new motor behaviour.

## MATERIALS AND METHODS

### Study Population

We included patients meeting the following criteria: (1)first-ever unilateral chronic (>6months) stroke; (2)Mini Mental Scale Examination(MMSE)[15]>24; (3)ability to perform 45° shoulder flexion while UL supported; (4)ability to be active for an hour; (5)no UL therapy during the study duration; (6)ability to understand the task and give written informed consent. Patients were excluded if they met any one of the followings: (1)ataxia/cerebellar stroke; (2)alcohol/drug abuse; (3)peripheral motor problems; (4)major psychiatric/other neurological disorders; (5)vision/hearing impairment; (6)neglect (Bells test)[16]; (7)shoulder pain/musculoskeletal impairment preventing passive ranging to the workspace; (8)<18 years old.

We screened 75 stroke survivors, and included 45 (supplementary figure 1). Patients were randomly allocated to one of three groups, according to the feedback given during adaptation [reward/punishment/neutral]. Randomization was stratified for age and time post-stroke. Fifteen healthy controls, matched to the neutral group for age and performing arm, were also included.

All participants gave written informed consent. The study was approved by the Joint Ethics Committee of the Institute of Neurology, UCL and the National Hospital for Neurology and Neurosurgery and was conducted in accordance with the Declaration of Helsinki.

### Experimental Task

We used a force-field (FF) adaptation paradigm[17]. Participants sat with their forehead supported in front of a workstation whilst holding the handle of a two-joint robotic manipulandum with their paretic arm. The forearm was stabilized by straps to a moulded cast and the trunk was belted to a chair. A horizontal mirror, 2cm above the hand, prevented direct vision of the arm, but showed a reflection of a screen mounted above. Visual feedback regarding hand position was provided by a white cursor (0.3cm diameter) continuously projected onto the screen (figure 1A).

The task consisted of centre-out fast ballistic movements to targets. Subjects had to initially bring the cursor within a 1-cm^2^ starting box in front of the body’s midline. Once the cursor was within the starting point, a white 1-cm^2^ target appeared 6cm from the starting position. Subjects were instructed that, when ready, they should make a fast, accurate, ‘shooting’ movement through the target, avoiding corrections. As the cursor crossed an imaginary 6cm radius circle centred at the starting position, a green endpoint dot appeared. After 500ms, the manipulandum returned the hand back to the start. For the main experiment, subjects were exposed to two targets. To encourage constant speed, the target turned red or blue if the movement was >500ms or <100ms, respectively (figure 1B).

For FF trials, the manipulandum produced a force proportional to the hand velocity. For a clockwise (CW) curl-field (pushing to the right) the force was:

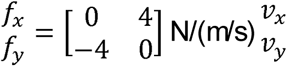

For counter-clockwise (CCW) curl-fields, the force direction was mirrored[17].

### Reward and Punishment Feedback

The reward group accumulated positive points, the punishment accumulated negative points and the neutral received no points. Points were calculated based on angular error as follows:*Reward:* 4 points: <1°; 3 points: 1-5°,2 points: 5-10°,1 point: 10-15°; 0 points:≥15°.*Punishment:*0 points: <1°;−1 points: 1-5°;−2 points: 5-10°;−3points:10-15°;−4 points: ≥15°,*Neutral.*Points were replaced by two zeros.

Both the points received on a trial-by-trial basis and the cumulative score of the block were shown (figure 1B). Subjects were informed that points had monetary value (3.57 pence/point) and depended on performance. The reward group started from £0 and earned money based on the accumulated points (£24.7±2), the punishment group was initially given £50 and lost money based on the cumulative negative points (average lost £24.5±1.7). The neutral group received £25 on day 3.

### Experimental Protocol

The same examiner tested all subjects across three consecutive days (figure 1C). Each session lasted around 2.5 hours. As sleep can enhance memories consolidation[18], participants were encouraged to sleep ≥6 hours every night, and sleep was assessed with a questionnaire[19].

### Day 1 (D1): Individual calibration of targets and perturbation

To ensure relatively accurate behaviour across patients and to select an error-enhancing force-field, on D1 participants familiarized with the task (six blocks, 80 trials each) with null trials (no FF) towards eight targets (25, 65, 115, 155, 205, 245, 295 or 335°CW from 0°, with 0° representing 12 on a clock). Based on performance, we selected for each subject the two targets in the same quadrant with the smallest average error and the FF direction (CW or CCW) enhancing this baseline error.

### Day 2 (D2): Adaptation under reward, punishment or neutral feedback

Participants were randomly allocated to the reward, punishment or neutral group (between-subject design) and performed 12 blocks (50 trials each) of reaching movements towards the two selected targets. Two baseline blocks were followed by 7 adaptation (CW or CCW FF) blocks, with reward/punishment/neutral feedback according to group allocation. In the washout (3 blocks), FF and reward/punishment/neutral feedback were removed to return performance to baseline. Participants were informed before beginning that they should expect the manipulandum to interfere with their performance, and that they should perform accurately whilst maintaining a constant speed. Short breaks were given after the second, fifth and tenth block. Participants could rest for a few minutes in between blocks if necessary.

### Day 3 (D3): readaptation at 24 hours

Participants were exposed to the same blocks as D2, but received neutral feedback.

**Figure 1.**
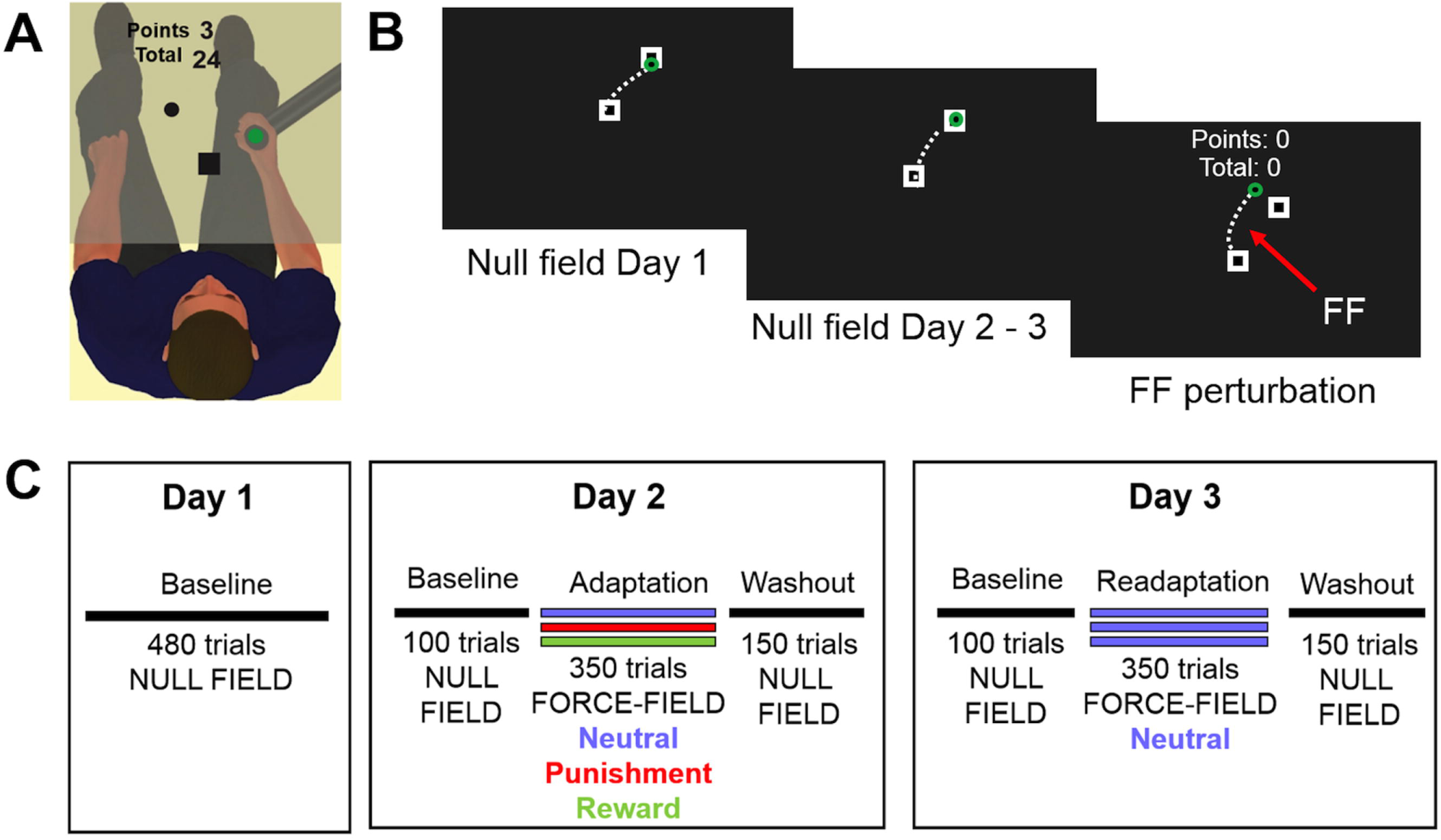
Task and protocol overview.**(A)** Experimental setup. **(B)** Experimental task. Participants moved the cursor from the starting point (central square) to a target on the screen. On Day 1, they had to reach towards one of eight targets, appearing in a pseudorandom order (null field day 1). On day 2 and 3, participants reached towards two selected targets which were chosen based on minimising baseline error (null field day 2 and 3). The perturbation consisted of a velocity-dependent force-field (red arrow) in the direction which increased baseline error (clockwise vs. counterclockwise). Reward and punishment feedback were represented by positive and negative points based on movement error, two uninformative zero instead of points appeared on the screen for the neutral group. **(C)** Experimental protocol. Participants were tested across three consecutive days. On day 1, they performed unperturbed reaching movements towards 8 targets (baseline: 6 blocks of 80 trials). Day 2 began with unperturbed reaching movements towards 2 targets (baseline: 2 block × 50 trials). This was followed by movements that were perturbed by a force-field (adaptation: 7 blocks × 50 trials) and in which participants received feedback according to their group (neutral, punishment or reward feedback). Finally, participants experienced another set of unperturbed trials (washout: 3 blocks × 50 trials). Day 3 was identical to day 2, except all groups received neutral feedback during the readaptation phase.

### Cognitive Tests and Functional Scales

To take into account possible confounding variables, subjects underwent a battery of validated tests and scales (supplementary table 1). The MMSE and the Bells test were used to assess eligibility. We then assessed executive functions, apathy, depression, fatigue, and sensitivity to reward and punishment. We used the Barthel Index to evaluate general functional level[20], and the Fugl-Meyer Assessment[21], the Modified Ashworth scale[6,22] and the Medical Research Council scale for strength[6,23] for the paretic UL. Handedness was evaluated using the Edinburgh Inventory[24]. At the end of each visit participants scored alertness and fatigue on a visual analogue scale.

### Data Collection and Analysis

The 2D (x,y) position of the hand was collected through custom C++ code (sampling rate=100 Hz). Data and statistical analysis were performed using Matlab (MathWorks, USA) and SPSS (IBM, USA). Movement onset was defined as the point at which velocity crossed 10% of peak velocity. Movement endpoint was defined as the position where the cursor breached the 6-cm target perimeter. To compare between subjects, errors of subjects receiving the CW FF were flipped.

Performance was quantified using angular error at peak velocity (AE_maxV_), i.e. the difference between the target angle and the angular hand position at the peak outward velocity(°). This has been used as a measure of feedforward control whilst excluding feedback processes[25]. To adjust for between-subjects baseline directional biases, AE_maxV_ on D2 and D3 were corrected by subtracting the average baseline AE_maxV_ of the corresponding day[26,27]. Reaction time (RT, time between target appearance and movement onset); movement time (MT, time between movement onset and movement end); peak velocity (MaxV); maximum velocity percentage (MaxV%, time point in movement when MaxV occurred); within subject variability (SD of AE_maxV_); and online corrections (difference between AE_maxV_ and angular endpoint error), were calculated for each trial. Trials in which angular error exceeded 60°[13] or MT or RT exceeded 1150ms (representing the mean +2.5 SD for both MT and RT) were removed (6.8% of trials). Epochs of all kinematics were created by averaging across 10 consecutive movements[27,28].

Difference between demographics, cognitive and functional scores were evaluated by one-way ANOVA (quantitative data) or Chi-square or Fisher exact test (proportions). Repeated-measures ANOVAs were used to compare MT, RT, MaxV, MaxV% and online corrections between groups (N, R, P) and phases (baseline, adaptation/readaptation, washout).

Due to the unfamiliarity with the manipulandum, unperturbed trials during D1 were also subject to a process of correction[29,30]. To evaluate this, we computed average sum of squared AE_maxV_during the first and last block of D1, and performed a repeated-measures ANOVA with group (N, R, P) and block (first, last). We used sum of square as we were interested in the absolute magnitude of the error, irrespective of direction.

To assess the amount of learning/adaptation independently from the co-contraction (i.e. stiffening) of the arm, we computed an adaptation index (AI):

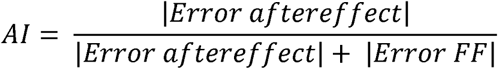

We considered as “aftereffect trials” the ones representing the initial error after the removal of the force-field. To select these, we performed an ANOVA across the average of every 2 trials for the first 10 trials (5 levels). On both days, we found a significant difference between trials 1-2 and 3-4 (D2: p=0.004; D3: p<0.001), and 3-4 and 5-6 (D2: p<0.001, D3: p=0.033). Based on this, we selected as “aftereffect trials” the first six trials after FF removal. Results were qualitatively similar by using an average between 2 and 6 trials. We defined as “force-field trials” the last block of the adaptation or readaptation. The AI could range zero, indicating no learning (but possibly co-contraction), to one, indicating complete learning[29]. This is based on the premise that learning is represented by zero error for FF trials but a large error in aftereffect trials (AI=1); no learning will lead to a large error in FF but zero error in the aftereffect trials (AI=0); and arm stiffening would cause zero error in both (AI=0).

To assess retention, i.e. the strength of the new motor memory, we calculated the average AE_maxV_across the last two washout blocks for D2 and D3 (AE_retention_)[13].

To account for differences in motor and cognitive functions, a principal component analysis (PCA) was conducted on the functional and cognitive scores, with varimax orthogonal rotation. The Kaiser-Meyer-Olkin (KMO=0.72) measure verified the sampling adequacy, and all KMO values were >0.6. Bartlett’s test indicated that correlations were sufficiently large (χ^2^_(45)_=136.36, p<0.001).

Three components had eigenvalue over Kaiser’s criterion of 1 and explained 71% of the variance. We interpreted the first component as the motor level, the second as the psychomotor level and the third as the cognitive level (supplementary table 2). We used these components as covariates in independent one-way ANCOVAs to compare groups for AI D2, AI D3, AE_retention_ D2 and AE_retention_D3.

To assess savings, i.e. the presence of faster readaptation when re-exposed to the same perturbation [31], we calculated an average AE_maxV_ for the first two perturbation blocks and performed a repeated measure ANOVA with groups (N, R, P) and days (D2, D3)[32].

No statistical methods were used to predetermine the sample size, but this is in line with similar studies on motor learning in stroke[3–8]. Data were tested for normality using the Shapiro-Wilk test. Homogeneity of variance was evaluated using Mauchly’s or Levene tests. When sphericity was violated, Greenhouse-Geisser (epsilon, ⍼<0.75)/Huynh-Feldt (⍼>0.75) corrections or Brown-Forsythe tests were used. Significance level was set at p<0.05. LSD post-hoc tests were conducted when warranted. Effect size was provided by partial eta (η^2^).

## RESULTS

### Demographics and Cognitive Functions were Similar between Groups

Demographic, and cognitive parameters were similar between groups (table 1& supplementary table 3). Healthy controls were similar to patients for age and main cognitive tests (supplementary table 4).

**Table 1.** Patient’s demographics and clinical characteristics

| | **N (n=15)** | **R (n=15)** | **P (n=15)** | $\lambda^2(2)$<br>*or F*<sub>(2,42)</sub> | **p** | **Effect size** |
| --- | --- | --- | --- | --- | --- | --- |
| **Sex (male)** | 9 (60) | 10 (66.7) | 7 (46.7) | 1.27 | 0.529 | 0.168 |
| **Age (years)** | 58.5±3.6 | 58.9±3.1 | 56.3±3.4 | 0.17 | 0.846 | 0.008 |
| **Education (years)** | 14±0.8 | 14.9±0.9 | 13.1±0.8 | 1.13 | 0.333 | 0.051 |
| **Stroke type (ischemic)** | 10 (66.7) | 13 (86.7) | 11 (73.3) | 3.21 | 0.66 | 0.267 |
| **Paretic limb (left)** | 9 (60) | 8 (53.3) | 9 (60) | 0.18 | 0.913 | 0.064 |
| **Dominant side affected** | 3 (20) | 6 (40) | 6 (40) | 1.8 | 0.407 | 0.200 |
| **Post-stroke (months)** | 58.3±13.2 | 41.5±5.4 | 45±13.7 | 0.6 | 0.552 | 0.028 |
| **Stroke site\*** |  |  |  |  |  |  |
| Cortical | 7 (46.7) | 9 (60) | 8 (53.3) | 0.07 | 0.966 | 0.046 |
| Subcortical | 3 (20) | 3 (20) | 3 (20) | 0.07 | 0.966 | 0.046 |
| **Functional scores** |  |  |  |  |  |  |
| FMA-UL | 41.8±3.4 | 49.8±3.3 | 45.6±3.5 | 1.39 | 0.26 | 0.062 |
| Barthel Index | 90.3±3.6 | 95±1.4 | 94±1.5 | 1.05 | 0.360 | 0.047 |
| Spasticity | 0.9±0.2 | 0.5±0.1 | 1±0.2 | 2.12 | 0.122 | 0.095 |
| Muscle strength | 2.8±0.4 | 3.8±0.4 | 3.4±0.3 | 2.42 | 0.101 | 0.103 |
| **Psychoactive drugs** |  |  |  |  |  |  |
| Antidepressants | 2 (13.3) | 1 (6.7) | 2 (13.3) | 0.45 | 0.799 | 0.1 |
| Antianxiety | 2 (13.3) | 0 (0) | 1 (6.7) | 2.14 | 0.762 | 0.218 |
| Muscle relaxants | 1 (6.7) | 1 (6.7) | 2 (13.3) | 0.55 | 1.000 | 0.110 |

Categorical values are indicated as number of patients (n) and the percentage this relates to in terms of each group (%), numeric values as mean ± SEM. Comparison between proportions is made with Chi-square test, comparison between means with one-way ANOVA test. Effect sizes are φ (phi) for chi-square test and η2 (eta squared) for one-way ANOVA. N=neutral; R=reward; P=punishment; FMA-UL=Fugl-Meyer Assessment Upper-Limb, measuring UL motor and sensory impairment. Scores range 0 to 66 with higher scores indicating better functioning; Barthel Index measures activities of daily living, scores range from 0 (totally dependent) to 100 (completely independent); Spasticity=averaged Modified Ashworth Scale (MAS) score from the shoulder, elbow and wrist joints. The MAS measures ranges 0 to 5, with higher scores indicating more spasticity; muscle strength=average Medical Research Council score measured from the shoulder flexors, elbow flexors and wrist extensor muscles, scores range 0 to 5, with higher scores indicating higher muscle strength. These muscles were chosen as they resist gravity in a reaching-out movement[33]. *Stroke site was not known in 12 patients (5 neutral, 3 reward and 4 punishment group).

### Day 1: Baseline Performance was Similar across Groups

MT, RT, MaxV, online corrections and variability on D1 were not significantly different across the patients groups (supplementary tables 5&6). The average sum of squared AE_maxV_ in the first and last block of D1 was different across blocks [*F*_(1,42)_=17.57, p<0.001, η^2^=0.295], but not between groups [*F*_(1,42)_=0.62, p=0.541, η^2^=0.029], with no group*block interaction [*F*_(2,42)_=0.695, p=0.505, η^2^=0.032]. This indicates similar baseline capability to correct for error across groups[29,30]. For each participant we then chose the target quadrant with the least amount of error, and the FF direction (CW vs. CCW) that enhanced this error (supplementary table 7).

### Day 2 and 3: Reward and Punishment Effects on Adaptation and Retention Kinematics, baseline performance and initial perturbation were similar across groups

Movement kinematics were similar across groups (supplementary tables 5&6). As expected, following target selection, variability was lower on D2 and D3 than D1 [*F* _(1,42)_=101.7, p<0.001, η^2^=0.708], but with no differences between groups [*F* _(2,42)_=0.34, p=0.715, η^2^=0.016] (supplementary table 8).

**Figure 2.**
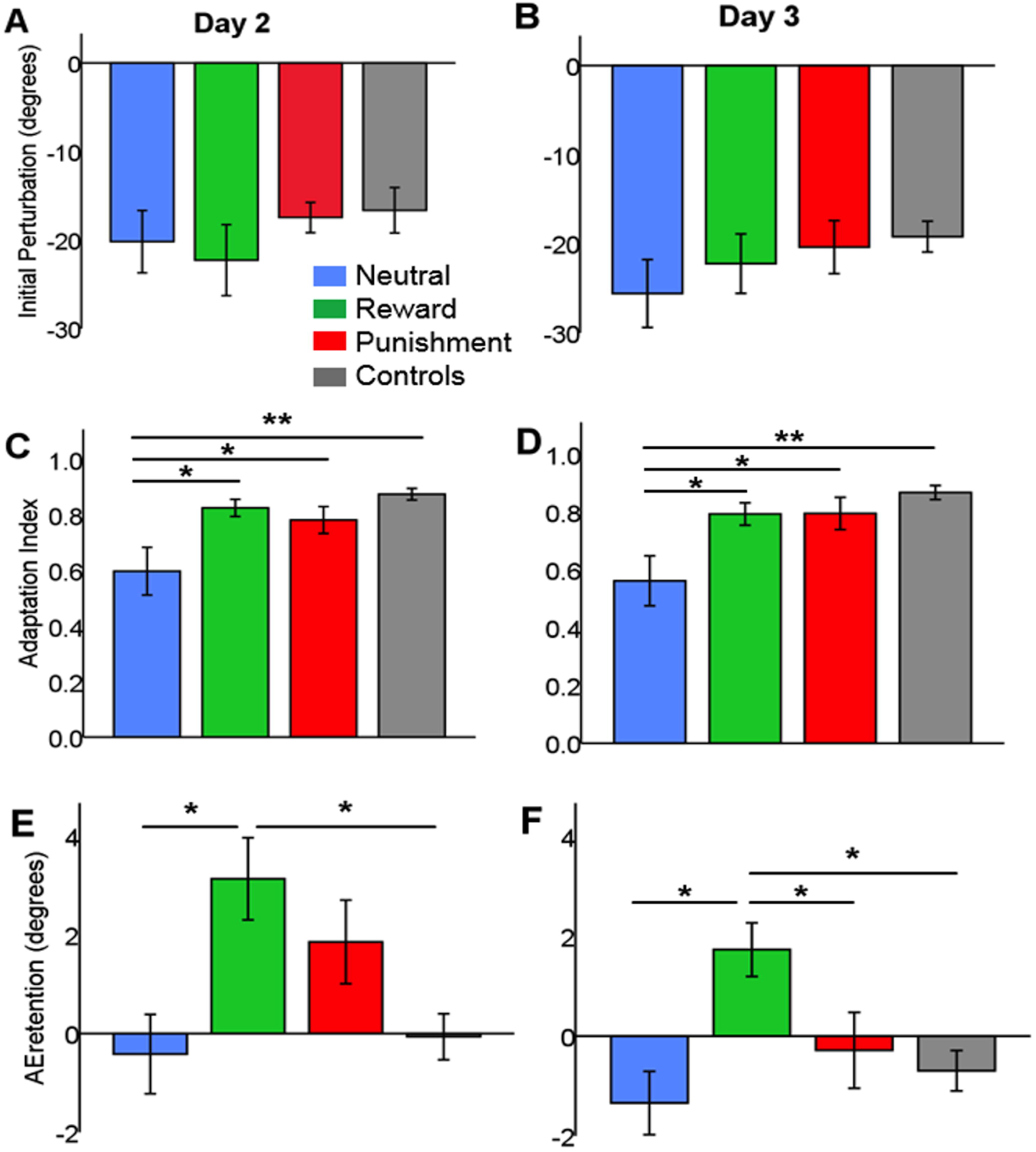
Average group data for day 2 (D2) and day 3 (D3). D2 and D3 angular error (degrees) at max velocity is shown during baseline, (re)adaptation and washout for the neutral stroke (blue), punishment stroke (red), reward stroke (green) and neutral healthy control (grey) groups. Values are mean (line) + SEM (shaded area) across epochs (average of 10 trials).

Baseline AE_maxV_ was similar across patients groups on both D2 [N:0.46+0.81°, R:-1.6+1°, P:-0.96+0.65°*F*_(2,42)_=1.5, p=0.235, η^2^=0.067] and D3 [N:0.13+0.67°, R:-0.2+0.74°, P:1.1+0.94°, *F*_(2,42)_=0.7, p=0.497, η^2^=0.033; figure 2]. The FF caused a similar initial perturbation across the three groups on both D2 [average AE_maxV_ across first two trials of FF, *F* _(2,42)_=0.5, p=0.577,η^2^=0.026; figure 3A] and D3 *F*_(2,42)_=0.6, p=0.551, η^2^=0.028; figure 3B].

**Figure 3.**
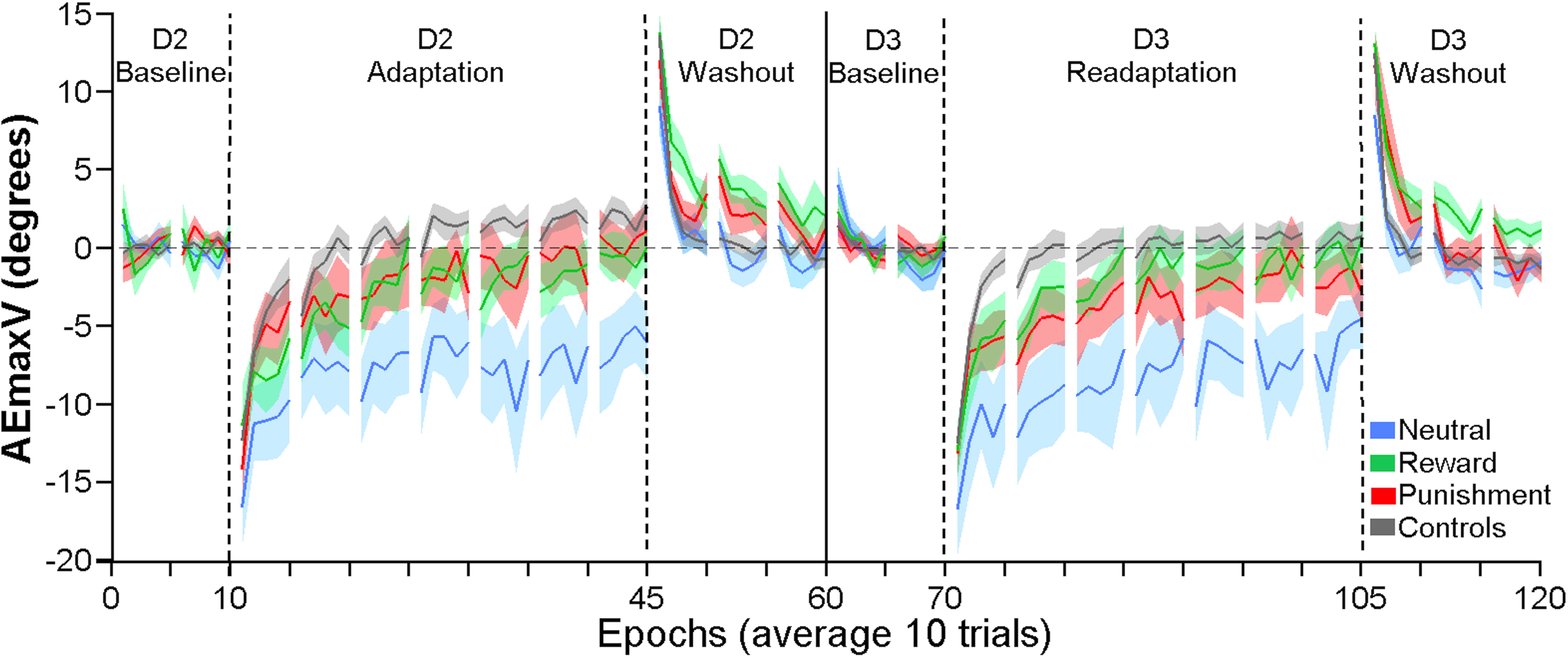
The initial perturbation (average angular error at peak velocity across the first two trials under force-field) was similar across groups on **(A)** day 2 (adaptation) and **(B)** Day 3 (readaptation). Adaptation index (AI) - ranging from 0 (no learning) to 1 (perfect learning) and evaluating learning independently from arm co-contraction day 2 (adaptation) was significantly lower in the neutral stroke group compared to the punishment stroke, the reward stroke and the neutral healthy controls groups. **(D)** AI on day 3 (readaptation) was significantly lower in the neutral stroke group relative to the punishment stroke, reward stroke, and neutral healthy control groups. **(E)** AE_retention_ on day 2-referring to the average angular error at peak velocity across last two washout blocks-was higher in the reward stroke group than in the neutral stroke group and the neutral healthy control group. No significant difference was found between the reward and punishment stroke groups.**(F)** AE_retention_ on day 3 was significantly higher in the reward stroke group versus the neutral stroke, punishment stroke and neutral healthy control groups. *p<0.05, **p<0.001.

### Reward and punishment were associated with greater adaptation and readaptation

Although all groups adapted, the reward and punishment group did to a greater extent (figure 2). After controlling for motor, psychomotor and cognitive functions, we found a significant effect of group on D2 AI [*F*_(2,39)_=3.422, p=0.043, η^2^=0.149; figure 3C], with lower adaptation in the neutral versus the reward (p=0.019) or punishment (p=0.050) groups.

Despite reward/punishment only being provided on D2, the improvements were maintained 24 hours later. Specifically, after controlling for the covariates, we found a main effect of group on D3 AI [*F*_(2,39)_=3.271, p=0.049, η^2^=0.144; figure 3D], once again with lower readaptation in neutral than either the reward (p=0.038) or punishment (p=0.029) groups.

### Reward was associated with higher retention

All groups displayed substantial aftereffects during washout on both D2 and D3 (figure 2), but the retention of this aftereffect was different across patient groups [D2 AE_retention_; *F*_(2,42)_= 3.425, p=0.043, η^2^=0.149; figure 3E], with the neutral retaining less than the reward group (p=0.016). Interestingly on D3 [*F*_(2,42)_=7.102, p=0.002, η^2^=0.267; figure 3F], the reward group displayed a greater amount of retention than either the neutral (p=0.001) or punishment (p=0.008) groups. No savings were observed across the groups, with no effect of group [*F*_(2,42)_=1.8, p=0.179, η^2^=0.079] nor day [*F*_(1,42)_=0.37, p=0.544, η^2^=0.009].

### Healthy Controls Adapted similarly to the Reward and Punishment groups but Retained Less

Although our focus was on patient groups, we also tested a group of age-matched healthy controls under neutral feedback. These showed less variability [overall variability: 6.1+0.2, *F*_(3,56)_= 7.17, p<0.001, η^2^=0.278] and faster RTs than patients but no differences in other kinematic parameters (supplementary tables 9&10), nor baseline AE_maxV_ [D2: *F*_(3,56)_=1.3, p=0.284, η^2^=0.065; D3: *F*_(3,56)_= 0.67, p= 0.575, η^2^=0.035] or initial perturbation [D2: *F*
_(3,56)_=0.7, p=0.556, η^2^= 0.036; D3: *F*_(3,56)_=0.82, p=0.485, η^2^=0.042].

Healthy controls adapted and readapted. Adaptation was significantly different across groups [D2 AI: Brown-Forsythe *F*_(3,28.5)_=5.3, p=0.005; figure 3C], with controls performing similar to the reward (p=0.51) and punishment (p=0.217) groups but significantly better than the neutral stroke (p<0.001) group. The same was observed for readaptation [D3 AI: Brown-Forsythe *F*_(3,33.2)_=5.6, p=0.003, figure 3D], with controls adapting more than the neutral (p<0.001), but similarly to the reward (p=0.353) and punishment (p=0.365) stroke groups.

Interestingly, despite adapting and readapting as the reward and punishment groups, controls retained less than the reward group (D2: p=0.004; D3: p=0.006). Controls showed no savings (p=0.174).

## DISCUSSION

We show for the first time that providing reward or punishment-based feedback to stroke patients during a motor adaptation task can bring their performances to the levels of healthy subjects of the same age range. More strikingly, reward increases the retention of the new motor behaviour to a level even higher than healthy subjects.

### Reward and Punishment Increased Learning

Although experiencing 350 trials, patients within the neutral group were unable to fully adapt. Remarkably, by providing reward or punishment, patients showed nearly full adaptation, similar to healthy controls. This was not explained by any differences in cognitive or functional scores between groups. Furthermore, day 1 performance was similar between patient groups, suggesting comparable baseline ability to correct for error. Finally, by individually tailoring the task on day 2 and 3, we further limited any possible influence of between-subject differences.

We previously showed in young healthy subjects that punishment led to faster adaptation, whereas reward caused greater retention[13]. We partially replicated these results, but found both punishment and reward associated with increased adaptation. One could argue that this effect may have been partially triggered by the knowledge of results provided by the feedback[34]. Nevertheless, our points system was unlikely to provide substantial amount of information in comparison to the visual feedback itself (i.e. 1 point represented a range of at least 5°). Secondly, patients’ sensitivity to feedback could be different to young healthy subjects. However, although aging is associated with reduced sensitivity to reward and punishment, the relative difference indicates an age-related hypersensitivity to reward[35]. Therefore, if we assume that younger adults’ greater sensitivity to punishment during adaptation represents the expected difference (loss aversion), then the stroke patients’ (older adults) results could demonstrate a hypersensitivity to reward. This also suggests that the specific effect of punishment on adaptation found in our previous work may be explained through loss aversion, rather than the hypothesised effect on cerebellar activity[13].

The improvements observed in the reward and punishment groups were maintained 24 hours later despite no further motivational feedback being provided. However, across all groups, we observed no savings. This is most likely due to the 250 washout trials and the 24 hour gap between adaptation blocks, both of which are known to significantly impair savings[36]. These results indicate that reward/punishment not only can enhance within-session adaptation in stroke patients, but, by making them learn better in the first place, could have long lasting benefits even when the feedback is no longer provided.

### Reward Increased Retention

Motor adaptation paradigms are already being implemented in some rehabilitation settings, such as gait rehabilitation[10]. Nevertheless the acquired motor behaviour is quickly forgotten, thus limiting the use of these paradigms in clinical practice. We found here that rewarding patients during adaptation increased retention. Most importantly, this effect was still present after 24 hours, with patients who had been rewarded retaining even more than controls. This is in line with previous evidences [11,13,14], and is a promising step toward the use of reward and motor learning paradigms in rehabilitation.

One caveat of using the after-effect as measure of retention is that this is influenced by the forgetting of what has been previously learnt (true retention), but also by simultaneous learning from movement errors[37]. Retention can be assessed using error-clamp trials[38], but these provide additional reward because patients are always successful in these trials. Nevertheless, the size and persistence of an aftereffect during washout trials with vision has been used numerous times as a proxy of retention[4,10].

### Implications

Clinically meaningful motor improvements in chronic stroke patients generally appear possible only with a large amount of contact hours[39]. Therefore, developing interventions that reduce the amount of hours required is crucial. This exploratory study highlights for the first time the potential of targeted motivational feedback as a tool to enhance the amount of learning and retention within and between sessions. Motor adaptation was used here as a model process, and further investigations on the effects of reward/punishment feedback over long-term training regimes are warranted. Robotic devices already in use in clinical rehabilitation could produce error-enhancing force-fields although improvements from robot-assisted therapy may not generalize to everyday life activities[40]. Therefore, how the improvements seen with motivational feedback could be administered within a setting where more practical behaviours are learnt remains a relevant question.

## Conclusions

We showed for the first time that reward and punishment enhance motor adaptation in stroke patients to similar level as controls. These improvements are maintained across 24 hours. Our findings suggest that the engagement of motivational processes during motor learning-based therapies could be a promising adjunct to rehabilitation. This will motivate further investigation about the long-term effects of motivational feedback, and thus avenues for translating these promising results into rehabilitation.

## Acknowledgements

We wish to thank you Ulrike Hammerbeck for her comments on the study design, and the Stroke Association and Different Strokes for their help with recruitment.

### Funding

This work was supported by a European Research Council Starter Grant (ActSelectContext, 260424 to SB) and Starter Grant (MotMotLearn, 637488 to JG).

## REFERENCES

Pollock A, BaerG, Campbell P,etalPhysical Rehabilitation Approaches for the Recovery of Function and Mobility After Stroke Major Update. Stroke 2014;45:e202–e202.

Kitago T, Krakauer JW. Motor learning principles for neurorehabilitation. In: Good MPB and DC, editor. Handbook of Clinical Neurology. Elsevier 2013:93–103.

Haaland KY, Harrington DL. Limb-sequencing deficits after left but not right hemisphere damage. Brain Cogn 1994;24:104–22.

Patton JL, Stoykov ME, Kovic M,et al Evaluation of robotic training forces that either enhance or reduce error in chronic hemiparetic stroke survivors. Exp Brain Res 2006;168:368–83.

Platz T, Denzler P, Kaden B,etal Motor learning after recovery from hemiparesis. Neuropsychologia 1994;32:1209–23.

Scheidt RA, Stoeckmann T. Reach adaptation and final position control amid environmental uncertainty after stroke. J Neurophysiol 2007;97:2824–36.

Takahashi CD, Reinkensmeyer DJ. Hemiparetic stroke impairs anticipatory control of arm movement. Exp Brain Res 2003;149:131–40.

Winstein CJ, Merians AS, Sullivan KJ. Motor learning after unilateral brain damage. Neuropsychologia 1999;37:975–87.

Chang WH, Kim Y-H., Robot-assisted Therapy in Stroke Rehabilitation. Stroke 2013;15:174–81.

Reisman DS, Wityk R, Silver K, etal Locomotor adaptation on a split-belt treadmill can improve walking symmetry post-stroke. Brain 2007;130:1861–72.

Abe M, Schambra H, Wassermann EM,etal Reward improves long-term retention of a motor memory through induction of offline memory gains. Curr Biol 2011;21:557–62.

Sugawara SK, Tanaka S, Okazaki S, etal Social rewards enhance offline improvements in motor skill. PloS One 2012;7:e48174.

Galea JM, Mallia E, Rothwell J,et alThe dissociable effects of punishment and reward on motor learning. Nat Neurosci 2015;18:597–602.

Goodman RN, Rietschel JC, Roy A, et al Increased reward in ankle robotics training enhances motor control and cortical efficiency in stroke. J Rehabil Res Dev 2014;51:213–27.

Folstein MF, Folstein SE, McHugh PR. “Mini-mental state”. A practical method for grading the cognitive state of patients for the clinician. J Psychiatr Res 1975;12:189–98.

Gauthier L, Dehaut F, Joanette Y. The Bells Test: A quantitative and qualitative test for visual neglect. Int J Clin Neuropsychol 1989;11:49–54.

Shadmehr R, Mussa-Ivaldi FA. Adaptive representation of dynamics during learning of a motor task. J Neurosci 1994;14:3208–24.

Al-Sharman A, Siengsukon CF. Sleep enhances learning of a functional motor task in young adults. Phys Ther 2013;93:1625–35.

Ellis BW, Johns MW, Lancaster R, etal The St. Mary’s Hospital sleep questionnaire: a study of reliability. Sleep 1981;4:93–7.

Mahoney FI, Barthel DW. Functional evaluation: the Barthel Index. Md State Med J 1965;14:61–5.

Fugl-Meyer AR, Jääskö L, Leyman I, et al The post-stroke hemiplegic patient. A method for evaluation of physical performance. Scand J Rehabil Med 1975;7:13–31.

Bohannon RW, Smith MB. Interrater reliability of a modified Ashworth scale of muscle spasticity. Phys Ther 1987;67:206–7.

Medical Research Council (Great Britain). Aids to the investigation of peripheral nerve injuries. London: H.M.S.O; 1975.

Oldfield RC. The assessment and analysis of handedness: the Edinburgh inventory. Neuropsychologia 1971;9:97–113.

Galea JM, Miall RC. Concurrent adaptation to opposing visual displacements during an alternating movement. Exp Brain Res 2006;175:676–88.

Ghilardi MF, Gordon J, Ghez C. Learning a visuomotor transformation in a local area of work space produces directional biases in other areas. J Neurophysiol 1995;73:2535–9.

Krakauer JW, Ghez C, Ghilardi MF. Adaptation to visuomotor transformations: consolidation, interference, and forgetting. J Neurosci 2005;25:473–8.

Galea JM, Vazquez A, Pasricha N,etal Dissociating the Roles of the Cerebellum and Motor Cortex during Adaptive Learning: The Motor Cortex Retains What the Cerebellum Learns. Cereb Cortex 2011;21:1761–70.

Smith MA, Shadmehr R. Intact ability to learn internal models of arm dynamics in Huntington’s disease but not cerebellar degeneration. J Neurophysiol 2005;93:2809–21.

van Beers RJ. Motor learning is optimally tuned to the properties of motor noise. Neuron 2009;63:406–17.

Kojima Y, Iwamoto Y, Yoshida K. Memory of learning facilitates saccadic adaptation in the monkey. J Neurosci 2004;24:7531–9.

Krakauer JW. Motor Learning and Consolidation: The Case of Visuomotor Rotation. Adv Exp Med Biol 2009;629:405–21.

Zackowski KM, Dromerick AW, Sahrmann SA,et alHow do strength, sensation, spasticity and joint individuation relate to the reaching deficits of people with chronic hemiparesis? Brain 2004;127:1035–46.

Vliet van PM, Wulf G. Extrinsic feedback for motor learning after stroke: what is the evidence? Disabil Rehabil 2006;28:831–40.

Bauer AS, Timpe JC, Edmonds EC,et alMyopia for the future or hypersensitivity to reward? Age-related changes in decision making on the Iowa Gambling Task. Emot 2013;13:19–24.

Criscimagna-Hemminger SE, Shadmehr R. Consolidation patterns of human motor memory. J Neurosci 2008;28:9610–8.

Hadipour-Niktarash A, Lee CK, Desmond JE,etalImpairment of retention but not acquisition of a visuomotor skill through time-dependent disruption of primary motor cortex. J Neurosci 2007;27:13413–9.

Scheidt RA, Reinkensmeyer DJ, Conditt MA,etalPersistence of motor adaptation during constrained, multi-joint, arm movements. J Neurophysiol 2000;84:853–62.

Cauraugh JH, Naik SK, Lodha N, et al Long-term rehabilitation for chronic stroke arm movements: a randomized controlled trial. Clin Rehabil 2011;25:1086–96.

Mehrholz J, Pohl M, Platz T, et alElectromechanical and robot-assisted arm training for improving activities of daily living, arm function, and arm muscle strength after stroke. Cochrane Database Syst Rev 2015;(11):CD006876.

